# Horseshoe crab genomes reveal the evolutionary fates of genes and microRNAs after three rounds (3R) of whole genome duplication

**DOI:** 10.1101/2020.04.16.045815

**Authors:** Wenyan Nong, Zhe Qu, Yiqian Li, Tom Barton-Owen, Annette Y.P. Wong, Ho Yin Yip, Hoi Ting Lee, Satya Narayana, Tobias Baril, Thomas Swale, Jianquan Cao, Ting Fung Chan, Hoi Shan Kwan, Ngai Sai Ming, Gianni Panagiotou, Pei-Yuan Qian, Jian-Wen Qiu, Kevin Y. Yip, Noraznawati Ismail, Siddhartha Pati, Akbar John, Stephen S. Tobe, William G. Bendena, Siu Gin Cheung, Alexander Hayward, Jerome H. L. Hui

## Abstract

Whole genome duplication (WGD) has occurred in relatively few sexually reproducing invertebrates. Consequently, the WGD that occurred in the common ancestor of horseshoe crabs ~135 million years ago provides a rare opportunity to decipher the evolutionary consequences of a duplicated invertebrate genome. Here, we present a high-quality genome assembly for the mangrove horseshoe crab *Carcinoscorpius rotundicauda* (1.7Gb, N50 = 90.2Mb, with 89.8% sequences anchored to 16 pseudomolecules, 2n = 32), and a resequenced genome of the tri-spine horseshoe crab *Tachypleus tridentatus* (1.7Gb, N50 = 109.7Mb). Analyses of gene families, microRNAs, and synteny show that horseshoe crabs have undergone three rounds (3R) of WGD, and that these WGD events are shared with spiders. Comparison of the genomes of *C. rotundicauda* and *T. tridentatus* populations from several geographic locations further elucidates the diverse fates of both coding and noncoding genes. Together, the present study represents a cornerstone for a better understanding of the consequences of invertebrate WGD events on evolutionary fates of genes and microRNAs at individual and population levels, and highlights the genetic diversity with practical values for breeding programs and conservation of horseshoe crabs.

## Background

Polyploidy provides new genetic raw material for evolutionary diversification, as gene duplication can lead to the evolution of new gene functions and regulatory networks (Holland 2003). Nevertheless, whole genome duplication (WGD) is a relatively rare occurrence in animals when compared to the fungi and plants (Van de Peer et al 2017). In animals, two rounds of ancient WGD occurred in the last common ancestor of the vertebrates, with additional rounds in some teleost fish lineages (Semon and Wolfe 2007; Jaillon et al 2009; Van de Peer et al 2017). Fixation of WGD or polyploidization has been considered a major force in shaping the evolutionarily success of vertebrate lineages by making fundamental changes in physiology and morphology, leading to the origin of new adaptations (Van de Peet et al 2009; Moriyama and Koshiba-Takeuchi 2018). Meanwhile, among the invertebrates, horseshoe crabs (Nossa et al 2014; Kenny et al 2016), spiders and scorpions (Schwager et al 2017) represent the only sexually reproducing invertebrate lineages that are known to have undergone WGD (Figure 1A).

**Figure 1.**
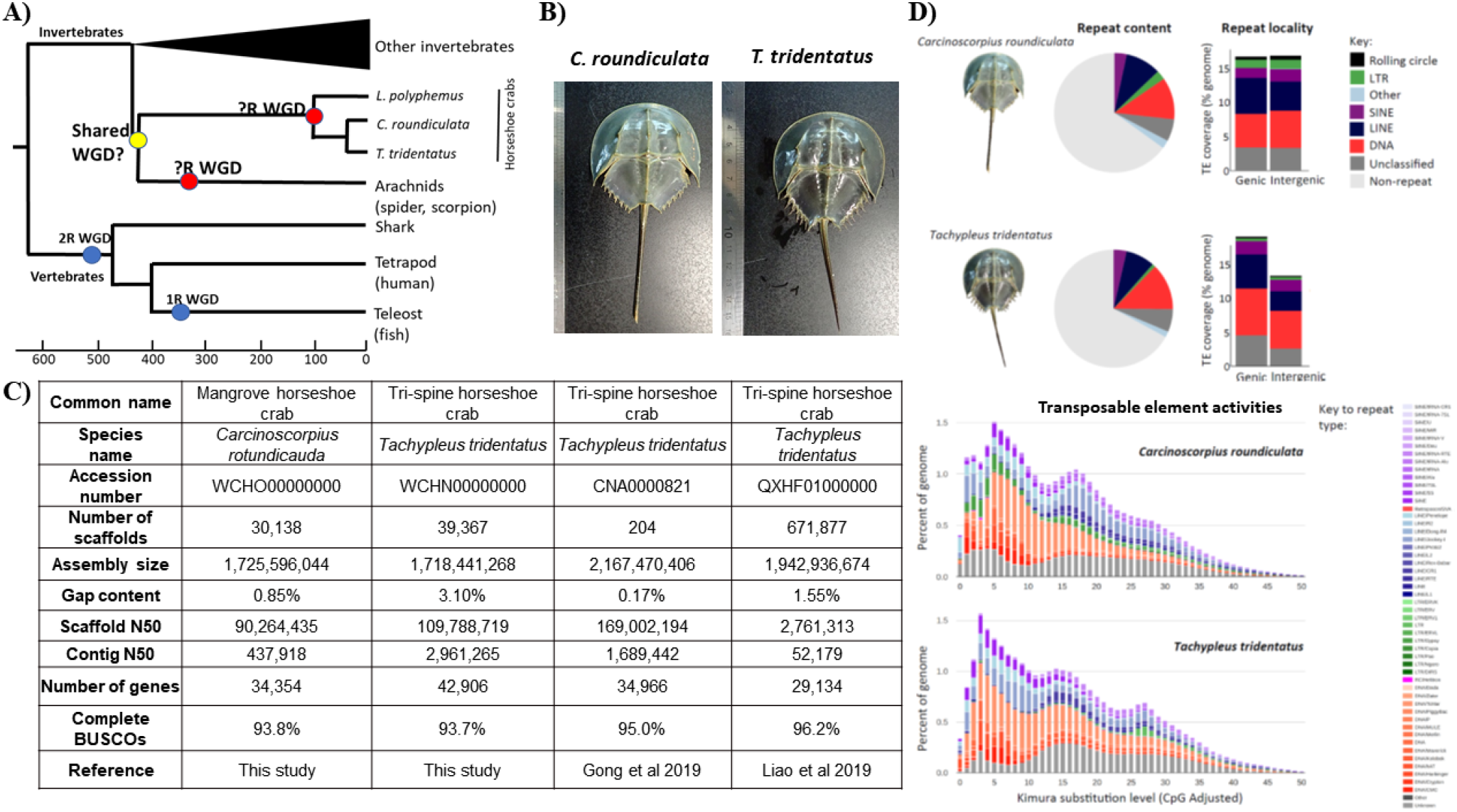
A) Schematic diagram illustrating the current knowledge of whole genome duplication (WGD) in animals. “?R” denotes unknown rounds of whole genome duplication; B) Pictures of horseshoe crabs *C. roundicultata* and *T. tridentatus;* C) Summary of genome assembly statistics of horseshoe crabs; D) Repeat content for the two horseshoe crab genomes, *C. rotundicauda* and *T. tridentatus:* Pie charts illustrating repeat content as a proportion of total genomic content; Repeat content present in genic verses intergenic regions; and Repeat landscape plots illustrating transposable element activity in each horseshoe crab genome.

Horseshoe crabs are considered to be ‘living fossils’, with the oldest fossils dated from the Ordovician period (~450 million years ago (Mya), Rudkin and Young 2009). However, despite this long history, there are only four extant species of horseshoe crabs worldwide: the Atlantic horseshoe crab (*Limulus polyphemus*) from the Atlantic East Coast of North America, and the mangrove horseshoe crab (*Carcinoscorpius rotundicauda*), the Indo-Pacific horseshoe crab (*Tachypleus gigas*), and the tri-spine horseshoe crab (*Tachypleus tridentatus*), from South and East Asia (John et al 2018). All extant horseshoe crabs are estimated to have diverged from a common ancestor that existed ~135 Mya (Obst et al 2012), and they share an ancestral WGD (Kenny et al 2016). A high-quality genome assembly was recently announced as a genomic resource for *T. tridentatus* (Gong et al 2019; Liao et al 2019), leaving an exciting research opportunity to analyse the genomes of other horseshoe crab species to understand how WGD reshapes the genome and rewires genetic regulatory network in invertebrates.

In the present study, we provide the first high quality genome of the mangrove horseshoe crab (C. *rotundicauda*), and a resequenced genome of tri-spine horseshoe crab (*T. tridentatus*). Importantly, we present evidence for the number of rounds of WGD that have occurred in these genomes, and investigate if these represent a shared event with spiders. We also examine the evolutionary fate of genes and microRNAs at both the individual and population level in these genomes. Collectively, this study highlights the evolutionary consequences of a unique invertebrate WGD, while also providing detailed genetic insights which will also be useful for various genomic, biomedical, and conservation measures.

## Results and Discussion

### High-quality genomes of two horseshoe crabs

Genomic DNA was extracted from single individuals of two species of horseshoe crab, *C. rotundicauda* and *T. tridentatus* (Figure 1B), and sequenced using Illumina short-read, 10X Genomics linked-read, and PacBio long-read sequencing platforms (Supplementary information S1, Table 1.1.1-1.1.2). Hi-C libraries were also constructed for both species sequenced using the Illumina platform (Supplementary information S1, Figure S1.1.1-1.1.2). For the final genome assemblies, both genomes were first assembled using short-reads, followed by scaffolding with Hi-C data. The *C. rotundicauda* genome assembly is 1.72 Gb with a scaffold N50 of 90.2 Mb (Figure 1C). The high physical contiguity of the genome is matched by high completeness, with 93.8% complete BUSCO core eukaryotic genes (Figure 1C). The *T. tridentatus* genome is 1.72 Gb with a scaffold N50 of 109.7 Mb and 93.7 % BUSCO completeness (Figure 1C). In total, the *C. rotundicauda* and *T. tridentatus* genome assemblies include 34,354 and 42,906 gene models, respectively. Furthermore, 89.8% of the sequences assembled for *C. rotundicauda* genome are contained on just 16 pseudomolecules, consistent with a near chromosome-level assembly (chromosome 2n=32, Iswasaki et al 1988, Supplementary information S1, Table 1.1.3).

To date, the only repeat data available for horseshoe crabs are two independent analyses of the tri-spine horseshoe crab *T. tridentatus*, which identified a repeat content of 34.61% (Gong et al 2019), and 39.96% (Liao et al 2019). In the present study, we provide the first analysis of repeat content in the genomes of different horseshoe crab species, by analysing repeats in our genome assembly for *T. tridentatus*, as well as our assembly for the mangrove horseshoe crab, *C. rotunicauda*. We find that repeat content is similar in both genomes, occupying approximately one third of total genomic content. Specifically, we identify a total repeat content of 32.99% for *T. tridentatus* and 35.01% for *C. rotunicauda*, of which the dominant repeats are DNA elements, followed by LINEs, with SINEs and LTR elements contributing just a small proportion of total repeat content (Figure 1D, Supplementary information S1, Table 1.2.1).

A large proportion of eukaryotic genomes is typically composed of repetitive DNA, and repeats are widely cited as being one of the key determinants of genome size (Chénais et al 2012). However, while the genome size for both species of horseshoe crab sequenced here is comparatively large for invertebrates, their repeat content is not unusually high (*C. rotundicauda:* 35.02%, *T. tridentatus:* 32.98%, Figure 1D, Supplementary information S1, Table 1.2.1). Instead, the comparatively large size of horseshoe crab genomes appears to be a consequence of multiple rounds of WGD, as discussed in greater detail below.

In the *C. rotundicauda* genome, repeats are evenly distributed across genic and intergenic regions (Figure 1D). However, in the *T. tridentatus* genome, a greater proportion of repeats are found in genic regions, due primarily to a higher density of DNA elements and LINEs, as well as unclassified elements (Figure 1D). Repeat landscape plots (Figure 1D) suggest a relatively similar pattern of historical transposable element activity for both horseshoe crab species. Recent activity appears to have tapered off more quickly in the *T. tridentatus* genome, particularly with respect to LTR elements and certain DNA elements (Figure 1D).

### Three rounds (3R) of whole genome duplications in horseshoe crabs

Initial efforts to analyse WGD in extant horseshoe crabs were from low-depth and genotyping-by-sequencing which hindered the understanding of WGD in these taxa (Nossa et al 2014; Kenny et al 2016). Recently, there have been two resequencing efforts for the horseshoe crab *T. tridentatus* (Gong et al 2019; Liao et al 2019), but our *T. tridentatus* genome assembly has the largest contig N50 (Figure 1C). Furthermore, our assembly for *C. rotundicauda* represents the first close to chromosomal-level genome assembly for this species. Consequently, the two high-quality horseshoe crab genomes presented in this study provide us with an unprecedented opportunity to address the issue of invertebrate WGD and its evolutionary consequences.

An important outstanding question is how many rounds of WGD occurred in the last common ancestor of horseshoe crabs, or alternatively if all rounds of WGD had occurred already in the ancestor of arachnids and horseshoe crabs (Figure 1A)? To address this question, we first investigated the number and genomic location of Hox cluster genes, which have played the role of a “Rosetta stone” for understanding animal evolution (Holland 2017). For example, the genome of the cephalochordate amphioxus contains only a single Hox gene cluster with 15 Hox genes, while the mouse genome contains four Hox gene clusters with 39 Hox genes, providing evidence that two rounds of WGD occurred between the most recent common ancestor of amphioxus and human (Putnam et al 2008; Holland 2013). In our horseshoe crab genomes for *C. rotundicauda* and *T. tridentatus*, the number of Hox genes was found to be 43 and 36, respectively (Figure 2A, Supplementary information S2). In *C. rotundicauda*, we found there are five Hox clusters, with other Hox genes located on additional small scaffolds; while in *T. tridentatus*, there are three Hox clusters, again with other Hox genes scattered across different scaffolds (Figure 2A). The situation is similar to the genome assembly of *L. polyphemus* (Nossa et al 2014), where our analyses showed that there are four Hox clusters with additional Hox genes located on different scaffolds. In a recent study of the *T. tridentatus* re-sequenced genome, the authors could only find two Hox clusters and could not identify the *Ftz* gene inside these clusters (Gong et al 2019). On contrary, our results suggested that there are three Hox clusters (including *Ftz*), and thus more than one round of WGD occurred in the lineage leading to extant horseshoe crabs.

**Figure 2.**
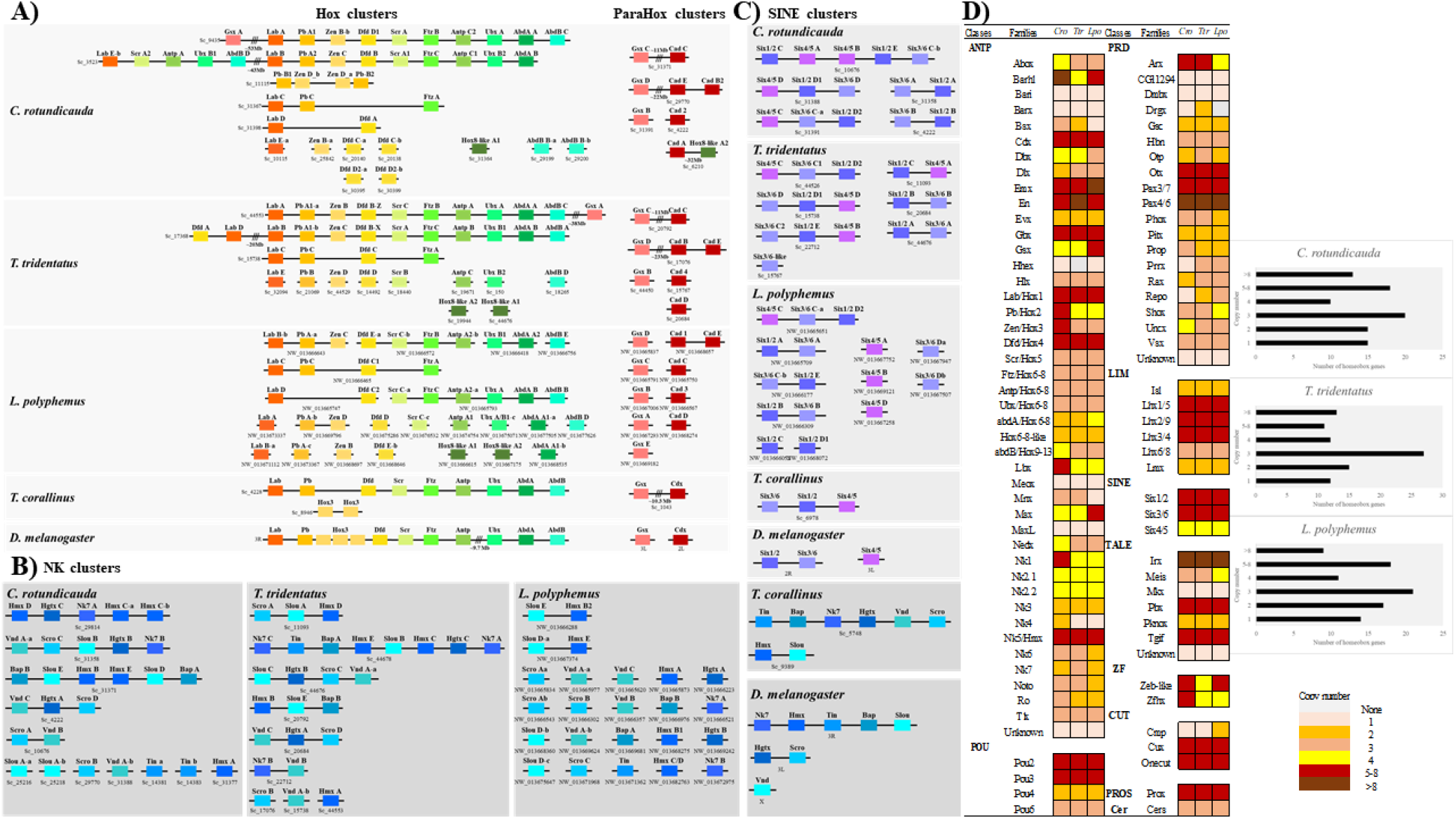
A) Genomic organisation of the Hox (left) and ParaHox (right) cluster genes in the horseshoe crab genomes. B) Genomic organisation of the NK and C) SINE cluster genes in the horseshoe crab genomes. D) Number of gene copies of homeobox genes in the horseshoe crab genomes.

We then investigated the sister cluster of the Hox genes – the ParaHox cluster genes, which are also highly clustered in bilaterians (Brooke et al 1998; Hui et al 2009; 2012). Similar to the Hox cluster genes, the invertebrate amphioxus contains only a single ParaHox gene cluster in its genome, while the ParaHox cluster genes are located on four chromosomes in human (Putnam et al 2008). In comparison, both the horseshoe crab genomes for *C. rotundicauda* and *T. tridentatus* contain two ParaHox clusters, composed of *Gsx* and *Cdx*, with other ParaHox genes located on three scaffolds. Meanwhile, in the genome assembly of *L. polyphemus* (Nossa et al 2014), perhaps due to the lower sequence continuity of the genome (i.e. low scaffold N50), only a single ParaHox cluster for *Cdx* was identified, with the other ParaHox genes were located on eight additional scaffolds (Figure 2A). In the situations relating to other well-known homeobox gene clusters, including the NK cluster and SINE clusters, as above, multiple clusters were revealed (Figure 2B-C). In *C. rotundicauda* and *T. tridentatus*, five and seven SINE clusters are found respectively, while in the genome assembly of *L. polyphemus* (Nossa et al 2014), four SINE clusters were revealed, with the other six genes located elsewhere in the genome.

Using genome-wide analyses of homeobox gene content in three horseshoe crab genomes, we find that many homeobox genes are present in more than 4 copies (Figure 2D, details are shown in Supplementary information S1, Table 1.2.2, Figure S1.2.1-1.2.5). These results suggest that at least two rounds (2R), and likely three rounds (3R) of WGD have occurred. The question then becomes, how many rounds of WGD have occurred. To address this question, we further carried out genome-wide synteny analyses to shed further light on the situation. As shown in Figure 3A, using a default of a minimum of 7 genes to define a syntenic block, most of the chromosomes of *C. rotundicauda* exhibit synteny with on other chromosomes, with most of them have a number between 4-8 (including its own copy). Thus, we propose that a 3R WGD occurred in the horseshoe crab.

**Figure 3.**
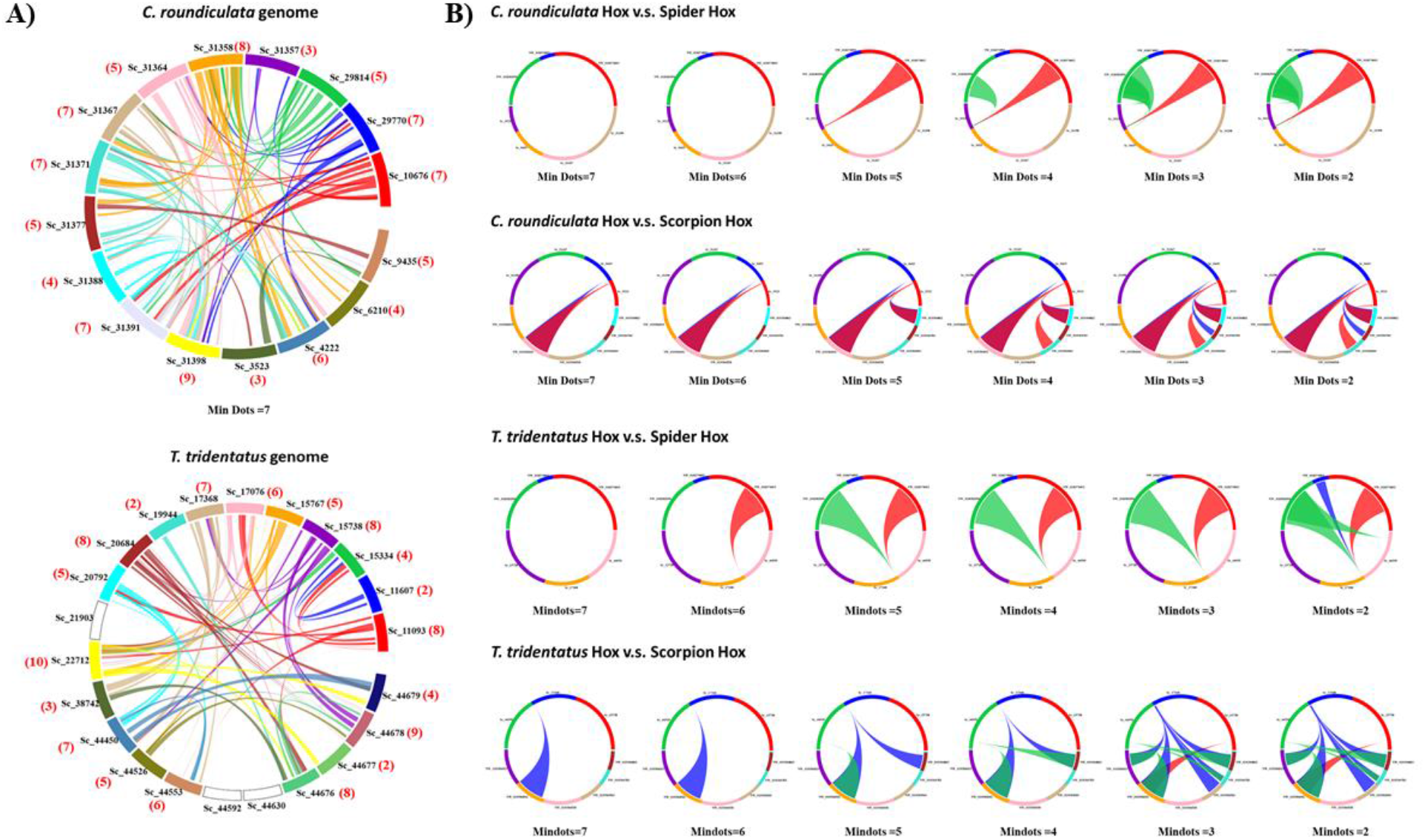
A) Synteny between different chromosomes of *C. roundiculata* and *T. tridentatus*. Note that the bracketed numbers highlighted in red refer to the numbers of chromosomes that syntenic blocks with that chromosome (counting include its own copy). B) Synteny relationships of Hox scaffolds of (Upper panel): *C. roundiculata*, spider and scorpion; (Lower panel) and *T. tridentatus*, spider, and scorpion.

### Shared or independent duplications with spider?

Another major unresolved question relating to horseshoe crab genomes is whether the reported cases of WGD in chelicerates constitute shared or independent events. Gene family analyses of spider and scorpion genomes have suggested that an ancient WGD is shared between them, independent of the WGDs that occurred in horseshoe crabs (Schwager et al 2017). Using the two horseshoe crab genome assemblies generated here, we tackled this important question from two different perspectives: (i) we performed analyses of synteny as a more rigorous examination of the question, and, (ii) we reconsidered recent evidence on phylogenetic relationships within the Chelicerata.

We first carried out the syntenic analyses between the Hox scaffolds of *C. rotundicauda* and the published spider and scorpion genomes (Schwager et al 2017) (Figure 3B). Despite no clear shared duplication event between *C. rotundicauda* and spider Hox, surprisingly, we observed syntenic relationships between two Hox scaffolds when using a minimum of 5 genes to define a syntenic block (Figure 3B). Similarly, in the syntenic comparison of Hox scaffolds of *T. tridentatus* and the published spider and scorpion genomes, we could observe syntenic relationships between two different Hox scaffolds when using a minimum of 5 genes to define a syntenic block (Figure 3B). In a less stringent condition of using a minimum of 2 genes to define a syntenic block, we additionally observed syntenic relationships between two other Hox scaffolds between *T. tridentatus* and spider (Figure 3B). Our data, suggested for the first time, that the WGD in horseshoe crab is a shared event with the WGD in spider and scorpion.

An important consideration necessary to fully understand WGD events identified from horseshoe crab genomes are the phylogenetic relationships between these animals. Horseshoe crabs have long been regarded as a monophyletic group (Xiphosura) and the sister group to the terrestrial chelicerate clade that includes spiders and scorpions (Arachinida). However, in a recent phylogenetic analysis using publicly available data, including three xiphosurans, two pycnogonids, and 34 arachnids, it has been suggested that the horseshoe crabs represent a group of marine arachnids (Ballesteros and Sharma 2019). On the other hand, another group of researchers recovered the Xiphosura as the sister group to the Arachnida (Lorano-Fernandez et al 2019), suggesting a single terrestrialisation event occurred after the last common ancestor of arachnids and horseshoe crabs diverged. Despite our data not being able to differentiate between these scenarios, we considered both situations while evaluating our data. In the Ballesteros and Sharma’s phylogeny, a shared WGD event occurred at the common ancestor of horseshoe crabs, spiders, and scorpions. On the other hand, the Loranzo-Fernandez et al phylogeny suggests that after the ancestral WGD at the ancestor of chelicerates and xiphosurans, massive gene losses may have happened in some lineages such as ticks and mites.

### Duplicated fates of noncoding microRNAs

With the availability of new transcriptomic data, especially the first small RNA transcriptomic data for both species of horseshoe crabs (Supplementary information 1, Table 1. 1.5-1.1.6), we analysed the evolutionary consequences of small noncoding RNAs after the WGD events in both *C. rotundicauda* and *T. tridentatus*. To reveal if duplicated microRNAs can also provide insights into the number of rounds of WGD, we first examined the number of paralogues for the bilaterian conserved set of 57 microRNAs, across three horseshoe crab genomes (Figure 4A). Of these microRNAs, 27 and 33 have more than 4 copies in *T. tridentatus* and *C. rotundicauda* respectively (Figure 4A). These data further support the hypothesis that 3R WGD occurred in the horseshoe crabs.

**Figure 4.**
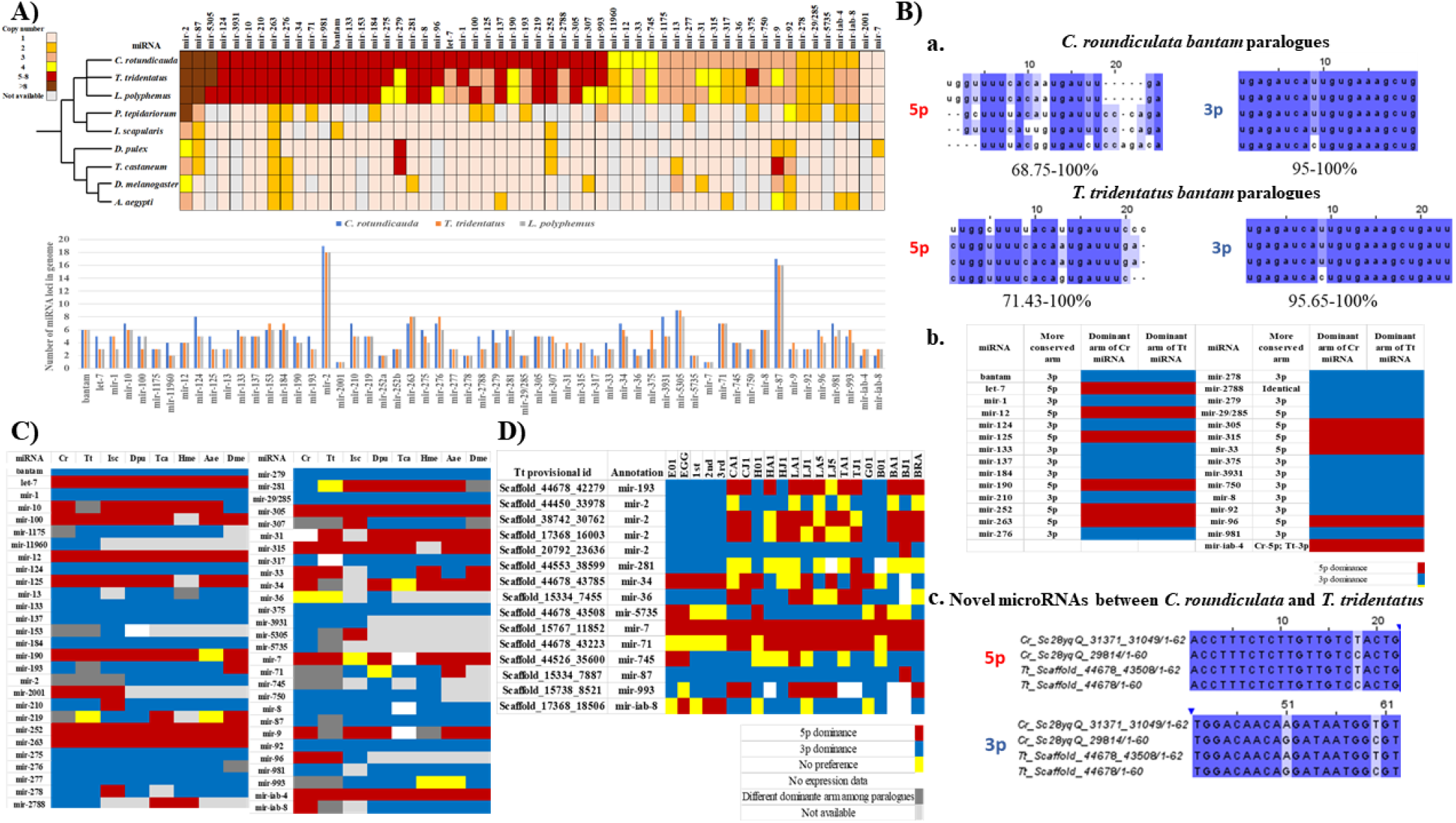
A) Number of gene copies of conserved microRNAs in the arthropod genomes. B) Sequence conservation and arm switching of horseshoe crab microRNAs. a) Degree of sequence conservation between bantam paralogues; b) Arm sequence conservation in relations to the dominant expression in between arms; c) Sequence alignment of novel microRNAs between the two horseshoe crabs. C) Comparison of microRNA arm preference among different arthropod species. Isc: *Ixodes scapularis*, Dpu: *Daphnia pulex*, Tca: *Tribolium castaneum*, Hme: *Heliconius melpomene*, Aae: *Aedes aegypti*, Dme: *Drosophila melanogaster*. D) MicroRNA arm switching cases among various tissue of Tt. Abbreviation: Egg-E01 and Egg; 1st, 2nd, 3rd instar-1st, 2nd, 3rd; Chelicerae-CA1, CJ1; Heart-H01, HA1, HJ1; 1st pair of leg-LA1, LJ1; 5th pair of leg-LA5, LJ5; Telson-TA1, TJ1; Gonad-G01; Blood-B01, BA1, BJ1; Brain: BRA; A-adult, J: juvenile. Arm preferrnce: blue-3p dominance, red-5p dominance, yellow-no preference, white-no expression.

To understand the fates of microRNA paralogues, we first analysed the sequence conservation/divergence of 41 conserved microRNA families and 4 chelicerate-specific microRNAs by aligning their sequences (Supplementary information S1, Figure S1.2.10). We found that the paralogues always have more sequence conservation in one arm (rather than showing similar conservation for both arms across paralogues) after WGD (Supplementary information S1, Figure S1.2.10). An example is illustrated for the microRNA bantam, where the sequence of the 5p arm is less conserved than the 3p arm between paralogues (Figure 4Ba).

To explore whether the more conserved microRNA arm correlates with expression level, we mapped small RNA reads to different paralogues. By eliminating microRNA species which have different arm usage between their paralogues or between horseshoe crab species, we found that, out of the 29 assessed microRNAs, 26 show a higher expression level/dominant arm usage at the conserved arm (Figure 4Bb, Supplementary information S3). For example, the 3p arm shows more sequence conservation between the bantam paralogues in horseshoe crabs, and their 3p arms also show higher expression levels than their 5p arms (Figure 4Bb, Supplementary information S3). The 26 conserved microRNAs identified as showing higher expression levels for the conserved arm serve as the first example correlating expression level and conservation of mature microRNA sequences in paralogues following WGD.

In addition to relatively old conserved microRNAs, we also investigated new/novel microRNAs which are specific to a certain horseshoe crab species, to understand the impact of WGD on these. A total of 12 novel microRNAs were identified and conserved between *C. rotundicauda* and *T. tridentatus* (Supplementary information S1, Figure S1.2.10). The identified novel microRNAs are highly conserved in sequences between orthologues than paralogues, an example is shown in Figure 4Bc, suggesting these horseshoe crab-specific novel microRNAs are born at the horseshoe crab ancestor after WGD.

In the common house spider *Parasteatoda tepidariorum* which is believed to have undergone a single round of WGD (Schwager et al 2017), paralogues of microRNAs were found to exhibit arm switching, a phenomenon whereby dominant microRNA arm usage is swapped among different tissues, developmental stages or species (Griffiths-Jones et al 2011; Leite et al 2016). We investigated microRNA arm switching in the sRNA transcriptomes generated here and compared this to their orthologues in various arthropods including fruitfly (*Drosophila melanogaster*), mosquito (*Aedes aegypti*), butterfly (*Heliconius melpomene*), beetle (*Tribolium castaneum*), water flea (*Daphinia pulex*), and tick (*Ixodes scapulari*) (Kozomara and Griffiths-Jones 2014; Fromm et al. 2020). By comparing dominant arm usage across different species, we found that many microRNAs, such as miR-2788, miR-281 and miR-iab-8 have undergone microRNA arm switching (Figure 4C, Supplementary information S3). Moreover, we also observed microRNA arm switching in cases of microRNAs throughout different developmental time or tissues (Figure 4D, Supplementary information S3). These findings are congruent with the spider microRNA study (Schwager et al 2017, Leite et al 2016).

In summary, the first investigation of microRNAs in horseshoe crabs provide another dimension for understanding the fates of duplicated noncoding microRNAs in invertebrates.

### WGD at population level

Another question that remains poorly explored is the evolutionary consequences of WGD on gene duplicates at the population level. Individuals of both *C. rotundicauda* and *T. tridentatus* were collected from different locations across Asia and subjected to genome sequencing (Figure 5A, Supplementary information S1, Table 1.2.3-1.2.4). As these genomes have undergone WGD, to confidently reveal their population structure, we only mapped sequencing reads to the mitochondrial genome and constructed the evolutionary trees from mitochondrial data. Distinct subpopulations can be identified within different regions in Asia, for example, the populations from Hong Kong formed a distinct group from other locations in Asia, which may be due to the strong ocean currents that had prevented the gene flows between these locations (Figure 5B; Supplementary information S1, Figure S1.2.6-1.2.7).

**Figure 5.**
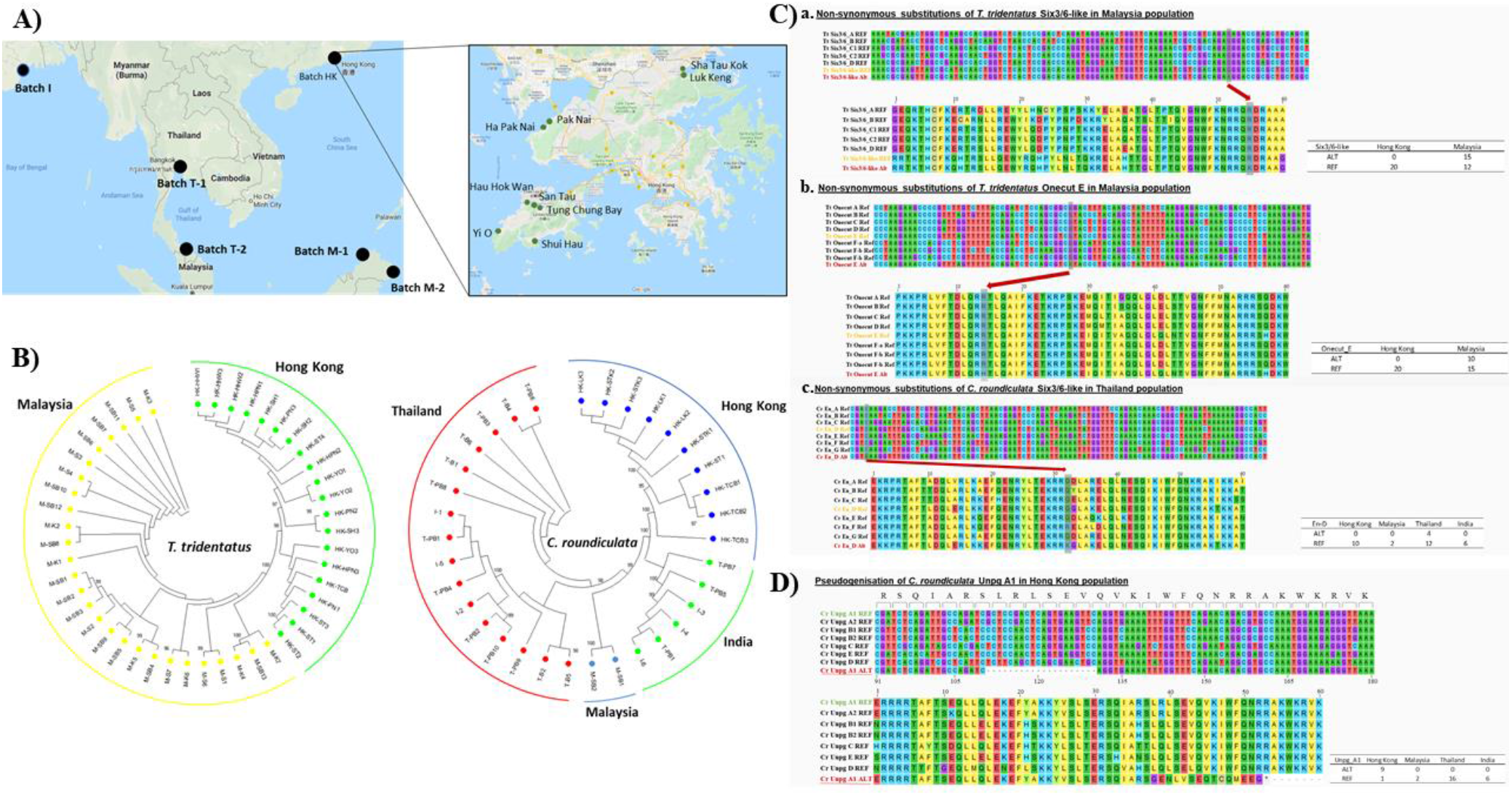
A) Geographical distribution of *C. roundicultata* and *T. tridentatus* collected samples; B) Phylogenetic trees of the collected samples. C) Non-synonymous substitutions of *T. tridentatus* (a) Six3/6-like; (b) Onecut-E genes in individuals collected in Malaysia; and (c). *C. roundiculata* En-D gene in individuals collected in Thailand population. D) Pseudogenisation of *C. roundiculata* Unpg-A1 gene in individuals collected in Hong Kong population.

Taking advantage of these population genomic data, we further asked the question of how dynamic the mutations at paralogues are in different individuals. With a focus on the homeodomain of the homeobox genes, we called single-nucleotide polymorphisms (SNPs) at the homeodomains of all annotated homeobox genes and found confident cases of both non-synonymous substitutions as well as pseudogenisation in the homeodomain of certain populations (Figure 5C; Supplementary information S4).

In *T. tridentatus*, non-synonymous substitutions at the homeodomain of Six3/6-like and Onecut-E genes were revealed in certain individuals from Malaysia populations (Figure 5Ca-b). Similarly for *C. roundiculata*, non-synonymous substitutions at the homeodomain of the *En-D* gene were also revealed in some individuals from populations in Thailand (Figure 5Cc). This is the first evidence showing that different gene duplicates after WGD in invertebrates are under different rates of mutation and selection at the individual level.

Importantly, unique pseudogenisation was discovered in the paralogue of *Unpg* in many individuals in the *C. rotundicauda* population located in Hong Kong (Figure 5D). In 9 out of the 10 individuals captured in Hong Kong for sequencing, we found that there is an alternative form (ALT), with a deletion in *Unpg-A1* (Figure 5D). Given that homeodomains are standardised as transcription factors with a sequence length of ~60-63 amino acids (Holland 2013), the deletion suggests that in these individuals these genes are in the process of becoming pseudogenes. This is the first evidence demonstrating the ongoing and dynamic mutation rate of paralogues at population level after WGD in invertebrates.

## Conclusion

WGD remains an understudied area, particularly in invertebrates such as the horseshoe crabs, despite its considerable importance in animal evolution. This study provides evidence of the 3R WGD events in horseshoe crabs, and sheds light on the evolutionary fates of genes and microRNAs at both the individual and population levels, as well as highlighting the genetic diversity of these amazing animals, with importance for understanding their evolution, genomics, and practical value for breeding programs and conservation.

## Materials and methods

### DNA, mRNA and sRNA extraction and sequencing

Genomic DNA of the horseshoe crabs *C. rotundicauda* and *T. tridentatus* was isolated from the leg muscle of a single individual in each case, using the PureLink Genomic DNA Kit (Invitrogen). In addition, different tissues were dissected and homogenized in Trizol reagent (Invitrogen), and total RNA was isolated following the manufacturers’ instructions.

Blood samples of both species of horseshoe crab were drawn by syringe and directly transferred into Trizol reagent for RNA extraction. For egg, 1^st^, 2^nd^ and 3^rd^ instars of *T. tridentatus*, whole individuals were used for RNA extraction. Extracted gDNA was subject to quality control using gel electrophoresis. Qualified samples were sent to Novogene and Dovetail Genomics for library preparation and sequencing. In addition, a Chicago library was prepared by Dovetail Genomics using the method described by Putnam et al (2016). Briefly, ~500ng of high molecular weight gDNA (mean fragment length = 55 kb) was reconstituted into artifical chromatin *in vitro* and fixed with formaldehyde. Fixed chromatin was digested with DpnII, the 5’ overhangs filled in with biotinylated nucleotides, and free blunt ends were ligated. After ligation, crosslinks were reversed, and the DNA purified. Purified DNA was treated to remove biotin that was not internal to ligated fragments. The DNA was then sheared to ~350 bp mean fragment size and sequencing libraries were generated using NEBNext Ultra enzymes and Illumina-compatible adapters. Biotin-containing fragments were isolated using streptavidin beads before PCR enrichment of each library. The libraries were sequenced on the Illumina HiSeq X platform. Dovetail HiC libraries were prepared as described previously (Lieberman-Aiden et al 2009). Briefly, for each library, chromatin was fixed with formaldehyde in the nucleus and then extracted Fixed chromatin was digested with DpnlI, the 5’ overhangs filled in with biotinylated nucleotides, and free blunt ends were ligated. After ligation, crosslinks were reversed and the DNA purified. Purified DNA was treated to remove biotin that was not internal to ligated fragments. The DNA was then sheared to ~350 bp mean fragment size and sequencing libraries were generated using NEBNext Ultra enzymes and Illumina-compatible adapters. Biotin-containing fragments were isolated using streptavidin beads before PCR enrichment of each library. Details of the sequencing data can be found in Supplementary information S1, Table 1.1.1-1.1.2.

Total RNA was subject to quality control using a Nanodrop spectrophotometer (Thermo Scientific), gel electrophoresis, and analysis using the Agilent 2100 Bioanalyzer (Agilent RNA 6000 Nano Kit). High quality samples underwent library construction and sequencing at Novogene; polyA-selected RNA-Sequencing libraries were prepared using TruSeq RNA Sample Prep Kit v2. Insert sizes and the concentration of final libraries were determined using an Agilent 2100 bioanalyzer instrument (Agilent DNA 1000 Reagents) and real-time quantitative PCR (TaqMan Probe) respectively. Small RNA (<200 nt) was isolated using the mirVana miRNA isolation kit (Ambion) according to the manufacturer’s instructions. Small RNA was dissolved in the elution buffer provided in the mirVana miRNA isolation kit (Thermo Fisher Scientific) and submitted to Novogene for HiSeq Small RNA library construction and 50 bp single-end (SE) sequencing. Detailed information for the sequencing data can be found in Supplementary information S1, Table 1.1.5-1.1.6.

### Genome, mRNA transcriptome, and sRNA assembly and annotation

To process the Illumina sequencing data, adapters were trimmed and reads were filtered using the following parameters “-n 0.1 (i.e. removal if N accounted for 10% or more of reads) −l 4 −q 0.5 (i.e. removal if the quality value is lower than 4 and accounts for 50% or more of reads)”. FastQC was run for quality control (Andrew 2010). If adapter contamination was identified, adapter sequences were removed using minion (Davis et al. 2013). Adapter trimming and quality trimming was then performed with cutadapt v1.10 (Martin 2011). For each species, k-mers of the Illumina PE library of 500 bp insert size were counted using DSK version 2.1.0 with k=25 (Rizk et al. 2013), and estimation of genome size, repeat content, and heterozygosity were analysed based on a k-mer-based statistical approach using the GenomeScope webtool (Vurture et al. 2017). Kraken was used to estimate the percentage of reads that may results from contamination from bacteria (Wood and Salzberg 2014). Chromium WGS reads were separately used to make a *de novo* assembly using Supernova (v 2.1.1), with the parameter “--maxreads=231545066” for *C. rotundicauda*, and “-- maxreads=100000000” for *T. tridentatus*, respectively. The *de novo* assembly, shotgun reads, Chicago library reads, and Dovetail HiC library reads were used as input data for HiRise, a software pipeline designed for using proximity ligation data to scaffold genome assemblies (Putnam et al, 2016). An iterative analysis was conducted. First, Shotgun and Chicago library sequences were aligned to the draft input assembly using a modified SNAP read mapper (http://snap.cs.berkeley.edu). The separation of Chicago read pairs mapped within draft scaffolds was analysed by HiRise to produce a likelihood model for genomic distance between read pairs, and the model was used to identify and break putative misjoins, to score prospective joins, and to make joins above a threshold. After aligning and scaffolding Chicago data, Dovetail HiC library sequences were aligned and scaffolded following the same method. After scaffolding, shotgun sequences were used to close gaps between contigs.

Raw sequencing reads of the transcriptomes were pre-processed with quality trimmed by trimmomatic (version 0.33, with parameters “ILLUMINACLIP:TruSeq3-PE.fa:2:30:10 SLIDINGWINDOW:4:5 LEADING:5 TRAILING:5 MINLEN:25”, Bolger et al. 2014). For the nuclear genomes, the genome sequences were cleaned and masked by Funannotate (v1.6.0, https://github.com/nextgenusfs/funannotate) (Palmer and Stajich 2018), the softmasked assembly were used to run “funannotate train” with parameters “ --stranded RF -- max_intronlen 350000” to align RNA-seq data, ran Trinity, and then ran PASA (Haas et al 2008). The PASA gene models were used to train Augustus in “funannotate predict” step following manufacturers recommended options for eukaryotic genomes (https://funannotate.readthedocs.io/en/latest/tutorials.html#non-fungal-genomes-higher-eukaryotes). Briefly, the gene models were predicted by funannotate predict with parameters “--repeats2evm --protein_evidence uniprot_sprot.fasta --genemark_mode ET -- busco_seed_species arthropoda --optimize_augustus --busco_db arthropoda --organism other --max_intronlen 350000”, the gene models predicted by several prediction sources including GeneMark (Lomsadze et al 2005), high-quality Augustus predictions (HiQ), PASA (Haas et al 2008), Augustus (Stanke et al 2006), GlimmerHMM (Majoros et al, 2003) and snap (Korf 2004) were passed to Evidence Modeler (Haas et al 2008) (EVM Weights: {‘GeneMark’: 1, ‘HiQ’: 2, ‘pasa’: 6, ‘proteins’: 1, ‘Augustus’: 1, ‘GlimmerHMM’: 1, ‘snap’: 1, ‘transcripts’: 1}) and generated the final annotation files, and then used of PASA (Haas et al 2008) to update the EVM consensus predictions, added UTR annotations and models for alternatively spliced isoforms. The protein-coding genes which cannot hit to nr db by DIAMOND blastp (version v0.9.22.123) (Buchfink B et al 2015) with evalue 1e-5 were removed.

To process small RNA data, we removed small RNA sequencing raw reads with Phred quality score less than 20, and adaptor sequences were trimmed. Processed reads of length 18bp to 27bp were then mapped to their respective horseshoe crab genome and analyzed using the mirDeep2 package (Friedlander et al 2011). To identify conserved microRNAs, the predicted horseshoe crab microRNA hairpins were compared against metazoan microRNA precursor sequences from miRBase (Kozomara and Griffiths-Jones 2014) using BLASTn (e value 0.01) (Altschul et al 1990). Predicted microRNAs which did not have significant sequence similarity to any of the microRNAs in miRBase were manually examined. Novel microRNAs were defined only when they fulfilled the unique features of microRNAs (Fromm et al 2020, MirGeneDB 2.0 https://mirgenedb.org/information). In addition, the copy number of microRNA loci was examined by using microRNA hairpins confirmed above to BLAST against each horseshoe crab genome.

### Annotation of repetitive elements

Repetitive elements were identified using an in-house pipeline. Firstly, elements were identified using RepeatMasker ver. 4.0.8 (Smit et al 2013) with the *Arthropoda* RepBase (Jurka et al 2005) repeat library. Low-complexity repeats were ignored (-nolow) and a sensitive (-s) search was performed. Following this, a *de novo* repeat library was constructed using RepeatModeler ver. 1.0.11 (Smit et al 2015), including RECON ver. 1.08 (Bao et al 2002) and RepeatScout ver. 1.0.5 (Price et al 2005). Novel repeats identified by RepeatModeler were analysed with a ‘BLAST, Extract, Extend’ process to characterise elements along their entire length (Platt et al 2016). Consensus sequences and classification information for each repeat family were generated. The resulting *de novo* repeat library was utilised to identify repetitive elements using RepeatMasker. Repetitive element association with genomic features were determined using BedTools ver. 2.26.0 (Quinlan et al 2010). “Genic” repetitive elements were defined as those overlapping loci annotated as genes ± 2kb and identified using the BedTools window function. All plots were generated using Rstudio ver. 1.2.1335 with R ver. 3.5.1 (Team 2013) and ggplot2 ver. 3.2.1 (Wickham 2016).

### Annotation of gene families and phylogenetic analyses

Potential gene family sequences were first retrieved from the two genomes using tBLASTn (Altschul et al 1990). Identity of each putatively identified gene was then tested by comparison to sequences in the NCBI nr database using BLASTx. For homeobox gene retrieval, sequences were also analysed using the BLAST function in HomeoDB. For phylogenetic analyses of gene families, DNA sequences were translated into amino acid sequences and aligned to other members of the gene family; gapped sites were removed from alignments and phylogenetic trees were constructed using MEGA.

### Synteny analyses

Synteny blocks were computed using SyMAP v4.2 (Synteny Mapping and Analysis Program) with default parameters except Min Dots from 2 to 7 (Minimum number of anchors required to define a syntenic block = 2-7) and “mask_all_but_genes = 1” to mask non-genic sequence (Soderlund et al 2011).

### Population genomic analyses

After quality control using FastQC (Andrews 2010), adaptors and low-quality bases were removed from the read ends using FASTP (Chen et al. 2018) with “-- qualified_quality_phred 30 --length_required 25” and other default parameters, followed by a second round of quality control using FastQC. The trimmed reads were mapped to the unmasked mitochondrion genome (NC_012574 of *T. tridentatus* and NC_019623 of *C. rotundicauda*) using bwa (version 0.7.12-r1039) with default parameters. The mapped reads were sorted usning SortSam of picard, and duplicated reads were removed using MarkDuplicates of picard. HaplotypeCaller from the Genome Analysis Toolkit GATK (version 4, https://gatk.broadinstitute.org/hc/en-us) was used to estimate the general variant calling file for each individual, and then combined by GenotypeGVCFs to a single variant calling file. Hard filtering of the SNP calls was carried out with Fisher strand bias (FS > 60.0), mapping quality MQ < 40.0, and thresholding by sequencing coverage based on minimum coverage (DP < 100) and maximum coverage (DP > 1,500). The SNPs were annotated with SnpEff (version 4.3T, http://snpeff.sourceforge.net/index.html)(Cingolani et al. 2012).

Filtered SNPs were used to generate population tree. The model-based software program STRUCTURE Version 2.3.4. 81 was used for population analysis. To determine most appropriate k value, burn-in Markov Chain Monte Carlo (MCMC) replication was set to 50,000 and data were collected over 1,00,000 MCMC replications in each run. Two independent runs were performed setting the number of population (k) from 2 to 10 using a model allowing for admixture and correlated allele frequencies. The basis of this kind of clustering method is the allocation of individual samples to k clusters. The k value was determined based on the rate of change in LnP(D) between successive k, stability of grouping pattern across two run and sample information about the material in supplementary file S1. Evolutionary divergence of within and between four different location horseshoe crab samples was performed using MEGA 7 (Molecular Evolutionary genetic analysis) following maximum composite likelihood model with 1000 bootstrap iterations of all samples. Principal coordinate analysis (PCoA) and UPGMA phylogenetic analysis was conducted to further assess the population subdivisions. PCoA was performed based on distance matrix using DARwin V.6.0.21 and UPGMA tree was constructed based on the simple matching dissimilarity (DARwin).

Trimmed reads were mapped to the homeodomain sequences using bwa (version 0.7.12-r1039) with default parameters. The mapped reads were sorted using SortSam of picard, and duplicated reads were removed using MarkDuplicates of picard. HaplotypeCaller from the Genome Analysis Toolkit GATK (version 4, https://gatk.broadinstitute.org/hc/en-us) was used to estimate the general variant calling file for each individual, and then combined by GenotypeGVCFs to a single variant calling file. Hard filtering of the SNP calls was carried out with Fisher strand bias (FS > 60.0), mapping quality (MQ < 40.0), QualByDepth (QD < 2.0), MappingQualityRankSumTest (MQRankSum < −12.5), ReadPosRankSumTest (ReadPosRankSum < 8.0) as https://gatkforums.broadinstitute.org/gatk/discussion/2806/howto-apply-hard-filters-to-a-call-set. The filtered out SNPs were then annotated with SnpEff (version 4.3T, http://snpeff.sourceforge.net/index.html(Cingolani et al. 2012). The missense mutation of the homeobox domain were manually checked with samtools tview.

### MicroRNA arm switching detection

The expression levels of 5p and 3p arms of microRNAs in the horseshoe crabs were calculated based on the number of sequencing reads mapped to the respective arm region in the predicted microRNA hairpin using bowtie/mirDeep2. The expression of different arms of microRNAs from different species were mapped according to previous method (Marco et al 2010) or referred to the data from MirGeneDB 2.0 (Fromm et al 2020). The arm usage ratio (AUR) of each microRNA was calculated using the formula AUR = 5p/(5p+3p), where 5p and 3p refer to the read counts of predicted 5p and 3p arms respectively. The AUR ranged from 0 to 1, with smaller values indicating the tendency of 3p preference and larger values indicating the tendency of 5p preference. 5p and 3p dominance was defined where AUR >0.7 and <0.3 respectively. No arm preference was defined when AUR ranged from 0.3 to 0.7. The overall arm preference (OAP) of each horseshoe crab microRNA was defined by evaluating their arm dominance in multiple tissue samples. If more than 70% of all tissue samples showed one type of arm dominance, then this type of arm dominance was defined as the OAP of this microRNA. Otherwise, no OAP was defined.

## List of abbreviations

3R: three rounds;
WGD: whole genome duplication;
AUR: arm usage ratio;
OAP: overall arm preference

## Acknowledgements

The authors thank Peter Holland for discussion, B. Desany for helping with the assembly of 10X Genomics data, and F. Cheung, R. Leung, W. Tong, W. Yiu, and H. Yu for collection of some of the RNA. This research was supported by the Hong Kong Research Grant Council GRF Grant 14103516; Environment and Conservation Fund Project 28/2017; Agriculture, Fisheries and Conservation Department of HKSAR Government, and the School of Life Sciences of The Chinese University of Hong Kong (JHLH).

## Figure legends and Supplementary information

Supplementary information S1. Supplementary data.

Supplementary information S2. Information of homeobox gene sequence and genomic locations.

Supplementary information S3. MicroRNA contents and arm usage of the two horseshoe crabs.

Supplementary information S4. SNPs at the homeodomains of the two horseshoe crabs.

